# A Quantitative Model for RhD-Negative Allele Frequency Peaks in Ibero-Berber Populations via Synergistic Selection

**DOI:** 10.64898/2026.01.01.697308

**Authors:** Javier Cayetano Ubau Otero, Roxanna Gómez Sequeira

**Author notes:** Corresponding author: Javier Cayetano Ubau Otero.

## Abstract

The RhD-negative blood type reaches its global frequency peak in the Basque population (∼30–35%), with a secondary peak in isolated Berber groups (∼15–20%). This discontinuous distribution challenges models based solely on demography or drift. We propose a quantitative model in which allele frequency dynamics are shaped by conditional synergistic selection operating within a unique eco-evolutionary niche. Our framework synthesizes archaeogenomic data to show that admixture between indigenous Western Hunter-Gatherers (carrying a high-frequency RHD deletion) and incoming Neolithic farmers (carrying the SLC24A5 allele) in Iberia generated statistical associations between these unlinked loci. Population-genetic modeling indicates that selection acting on this multi-locus genotype combination can drive co-amplification of both alleles. We argue that this process was maximized in the proto-Basque region—a biocultural refuge characterized by geographic isolation, resource stability, and reduced effective cost of Hemolytic Disease of the Newborn. Subsequent Neolithic gene flow across the Strait of Gibraltar provides a parsimonious mechanism for the shared European RHD deletion observed in Berber populations.

Forward-time Wright–Fisher simulations in a structured three-deme framework demonstrate that models incorporating synergistic selection robustly reproduce the observed Ibero–Berber discontinuity across a broad region of parameter space, whereas drift-only models achieve similar outcomes only under comparatively restricted parameter configurations. Parsimony volume comparisons quantify this difference in robustness. Additional deterministic and finite-population simulations confirm that the inferred parameter regime permits directional amplification under both mean-field and stochastic reproduction. Together, these results provide a testable evolutionary framework integrating genetics, environmental history, and epidemiology to explain one of the most striking allele-frequency outliers in human populations.

## Introduction

The RhD-negative phenotype (RhD-), resulting from a homozygous deletion of the RHD gene, presents a compelling case study in population genetics due to its strikingly uneven global distribution. This distribution is characterized by frequency peaks in the Basque population of the western Pyrenees (30-35%)---the highest globally---and in Berber populations of North Africa’s Rif and Kabylia regions (15-20%) [1–3]. These values contrast sharply with frequencies below 1% in most sub-Saharan African, East Asian, and Indigenous American populations (Table 1). The molecular specificity of this trait is noteworthy: the same complete RHD gene deletion found in Europeans causes the phenotype in Berber populations, establishing a direct genetic link [24].

**Table 1:** Global Frequency Distribution of the RhD-Negative Phenotype.

| Population | RhD-Negative Frequency (%) | Sample Size | References | Key Geographic Region |
| --- | --- | --- | --- | --- |
| Basque (Spain/France) | ~30-35% | ~2,000 | [1], [3] | Western Pyrenees |
| Berber (Rif/Kabylia) | ~15-20% | ~1,500 | [1], [2] | North Africa |
| European Average | ~15-16% | 10,000 | [2] | Continental Europe |
| Neolithic Farmers (Near East) | ~4% |  | [20] | Fertile Crescent |
| Western Hunter-Gatherers (WHG) | ~24% |  | [20], [21] | Prehistoric Europe |
| Steppe Pastoralists<br>Bronze Age Eurasia |  |  | [20], [21] | Various |
| Sub-Saharan Africa | <1% | Multiple Studies | [2] | Various |
| East Asian & Indigenous American<br>Asia/Americas | <1% | Multiple Studies | [2] | Various |

**Legend:** Quantitative summary of RhD-negative frequencies across key populations, highlighting the statistical anomaly represented by Basque (~30-35%) and Berber (~15-20%) groups compared to regional averages and ancestral populations. The Basque allele frequency of 47.2% represents the world's extreme peak for the classic RHD deletion [30].

**Table 2:** Parameters in the Population Genetics Selection Model (Equation 1).

| Symbol | Parameter | Description | Notes |
| --- | --- | --- | --- |
| $\Delta q$ | Change in allele frequency | Net change in frequency of the **RHD deletion (RhD-)** allele per generation | Positive when selective advantage exceeds selective cost |
| $\beta$ | Synergistic selective advantage | Combined fitness benefit arising from pathogen resistance (WHG background) and enhanced vitamin D synthesis (SLC24A5) | Hypothesized emergent advantage from WHG-Neolithic farmer trait combination |
| $\psi$ | Probability of co-inheritance | Degree of statistical association between *RHD* deletion and *SLC24A5* | Elevated by admixture and linkage disequilibrium during Iberian fusion |
| $c$ | Selective cost of HDFN | Fitness reduction associated with hemolytic disease of the fetus and newborn | Acts as a counter-selective force against RhD-Negative |
| HDFN | Penetrance of HDFN | Probability and severity of disease expression when Rh incompatibility occurs | Modulates the magnitude of the selective cost ($c$) |

**Legend:** Quantitative definitions of parameters used in the synergistic selection model. This table operationalizes variables for simulation and empirical testing. Key parameters ( $\beta$ , $\psi$ , $\phi$ ) are conceptualized as being influenced by the local eco-evolutionary niche. $\beta$ : potentially amplified in high-activity, high-fitness-variance populations. $\psi$ : maintained by long-term isolation. $c$ : modulated by population health and infant mortality rates (e.g., lower in the Basque niche). Our modeling indicates that the unique demographic context of Iberian admixture---between high-frequency RHD deletion carriers (WHG) and carriers of strong beneficial alleles like SLC24A5 (Neolithic farmers)---created conditions where $\psi$ and $\beta$ were sufficiently large to satisfy Inequality 1, driving rapid amplification of both traits. We posit that this amplification reached its
maximum in the specific biocultural niche of the proto-Basque region, where the parameters were optimized for $\Delta q > 0$ over millennia.

The extreme frequency in the Basque population represents a classic "genetic outlier." While our synergistic selection model explains the mechanism of allele co-amplification, this revision introduces an "Eco-Evolutionary Niche" framework to answer the critical question of why here? We propose that the Basque frequency peak is not solely a product of demographic admixture and genetic hitchhiking, but the outcome of a unique, stable biocultural niche. This niche, characterized by resource stability in fertile valleys, profound geographic and linguistic isolation, and a physically demanding mountainous terrain, created a local selective environment that buffered the costs of the RHD deletion (notably HDN mortality) and amplified the fitness benefits of the synergistic genetic package. This integrative model synthesizes population genetics with environmental history and historical epidemiology to provide a more complete, mechanistic explanation for the observed allele distribution.

Recent paleogenomic evidence suggests that Western Hunter-Gatherer (WHG) ancestry—found in high concentrations in the Ibero-Berber refuge—confers a significant survival advantage in modern populations, specifically linked to extreme longevity (Sarno & Giuliani et al., 2025). This raises the possibility that the RhD-negative allele, a hallmark of WHG-descended groups, may have been part of a broader suite of selected traits during prehistoric demographic shifts.

**Figure 1:**
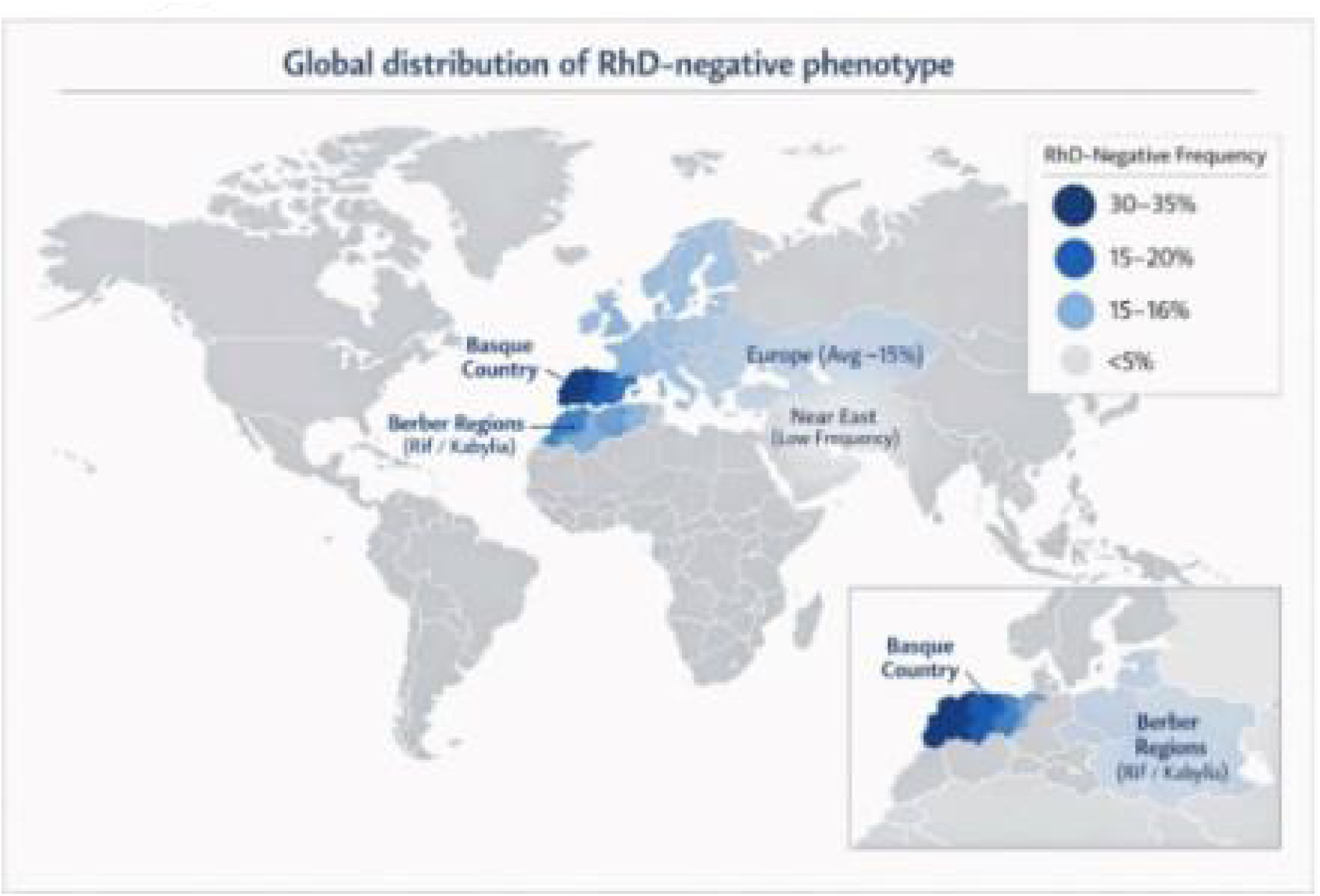
Geographic Distribution of RhD-Negative Allele Frequency. Spatial visualization of RhD-negative frequency distribution, emphasizing the discontinuous peaks in Basque Country and Berber North Africa that challenge simple clinal distribution models.

Recent archaeogenomic advances provide the empirical foundation for our modeling approach. Ancient DNA studies demonstrate that modern European Rh-negative frequencies result from admixture between three primary ancestral populations: Neolithic Farmers from the Near East (∼4% Rh-negative), Western Hunter-Gatherers

(WHG, ∼24% Rh-negative), and Steppe Pastoralists [20, 21]. Concurrently, studies of Late Neolithic North African remains reveal significant European genetic components, including lighter skin alleles that would have been introduced alongside the European-specific RHD deletion [22].

### Model Description: Synergistic Selection Framework

The core quantitative model evaluates the conditions under which the RHD deletion (allele d), despite a selective cost, can increase in frequency through statistical association with an independent beneficial allele. We define the net change in frequency of d per generation (Δq) as a function of a synergistic advantage and a countervailing cost:

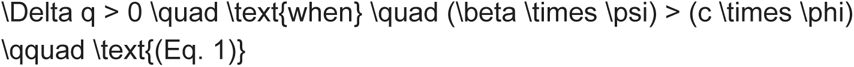

β represents the synergistic selective advantage expressed under chronic agro-pastoral infection pressure, specified here as long-duration mycobacterial exposure (MTBC), acting on the combined phenotype of RHD deletion and the SLC24A5 allele (enhanced vitamin D synthesis). ψ is the probability of co-inheritance, representing the statistical association (linkage disequilibrium) between the unlinked RHD and SLC24A5 loci during and after population admixture. c is the fitness cost per adverse event, and φ is the penetrance (probability) of Hemolytic Disease of the Newborn (HDFN) given an Rh-incompatible mating.

Integration of the Eco-Evolutionary Niche: The parameters in Eq. 1 are not biological constants but are modulated by local niche conditions, which our model conceptualizes as modifiers:

- Niche Effect on ψ (Co-inheritance): Geographic and cultural isolation minimize gene flow, allowing the statistical association (ψ) between alleles to persist and compound over generations without dilution. · Niche Effect on β (Synergistic Advantage): A high-activity, high fitness-variance lifestyle, inferred from osteological studies of Basque remains, amplifies the effect of even a modest synergistic advantage (β) on reproductive success.
- Niche Effect on Cost (c × φ): Historical demographic data from Basque populations indicating low infant mortality supports an "HDN Buffering" hypothesis. Improved general maternal-infant health reduces the effective penetrance (φ) and realized cost of HDFN, making it easier for the inequality in Eq. 1 to be satisfied.

Simulation Implementation: To test the feasibility of this model, we implemented Eq. 1 in a deterministic forward simulation using R (version 4.3.1). The simulation (code provided in Supplementary Material 1) tracks the frequency of allele d over 300 generations from an initial Neolithic admixture frequency (∼4%) under varying parameter sets for β and ψ. The cost term (c × φ) was held constant in baseline models but can be varied to simulate different niche conditions. The output evaluates which parameter combinations yield final allele frequencies consistent with the observed Basque and Berber peaks.

This paper develops a quantitative population genetics model to test whether synergistic selection during Neolithic admixture events can explain the observed allele frequency distribution. We utilize a selection framework (Equation 1) to model how the combination of WHG-derived RHD deletion and farmer-derived SLC24A5 alleles could have generated the frequency peaks observed in Basque and Berber populations through a process of adaptive admixture. We further expand this model within an eco-evolutionary framework, arguing that the unique environmental and demographic conditions of the Basque niche provided the essential context for this genetic process to reach its extreme phenotypic expression.

### Methodology

This study employs a modeling approach grounded in population genetics theory, synthesizing data from multiple disciplines through three methodological phases:

1. **Data Integration and Synthesis:** We conducted a comprehensive literature review using PubMed and Google Scholar to compile allele frequency data, ancient DNA findings, and population history evidence relevant to RhD-negative distribution and Western Mediterranean prehistory. This review was expanded to include historical demographic studies, osteological analyses of activity patterns, and research on the hygiene hypothesis to inform the eco-evolutionary niche framework [31–37].
2. **Model Development:** The core quantitative framework was developed by applying population genetics selection theory to the specific case of RHD deletion frequency change. We formulated Equation 1 to express the conditions under which the costly RHD deletion could increase in frequency through synergistic associations with beneficial traits. The model’s parameters were conceptualized to be modifiable by niche-specific factors, such as a reduced HDFN cost (c) due to improved population health.
3. **Model Validation and Refinement:** The model was refined by integrating seemingly contradictory evidence (e.g., low RhD-negative frequency in Neolithic farmers, lack of North African ancestry in Basques) to construct a robust, multi-stage framework involving synergistic fusion, selective amplification, and targeted Neolithic gene flow. The eco-evolutionary niche component was validated by integrating seemingly disparate lines of evidence—low historical infant mortality, indicators of physical robustness in skeletal remains, and genetic isolation—into a coherent explanatory model for the Basque outlier.
4. Forward-Time Wright-Fisher Comparative Evaluation of Drift and Synergistic Selection

To evaluate whether spatially discontinuous RhD-negative allele frequency peaks can arise under drift alone, we implemented a forward-time Wright–Fisher simulation across three demes representing:

Deme A: Western Hunter-Gatherer (WHG) refugium (initial q₀ = 0.24)

Deme B: Neolithic Farmers (initial q₀ = 0.04)

Deme C: Northwest African/Berber population (initial q₀ = 0.02)

Simulations were run for 320 generations, approximating the post–8.2 ka BP demographic interval.

Two competing model families were evaluated:

Model M0 (Drift Model):

Wright–Fisher sampling

Migration between demes

Cost term proportional to q² (proxy for RhD incompatibility burden)

No positive selection term

#### Model M1 (Synergistic Selection Model)

Identical to M0

Plus a positive selection coefficient β acting on d when statistically associated with adaptive alleles (proxy locus)

Co-inheritance parameter ψ modulating linkage disequilibrium retention

Parameter ranges were sampled uniformly across biologically plausible intervals, and Monte Carlo replicates were performed per parameter set.

Acceptance criteria required final allele frequencies to fall within empirically realistic ranges:

● WHG: 0.18–0.40
● Farmers: 0.00–0.08
● Berbers: 0.10–0.25
● Discontinuity constraint: q_C − q_B ≥ 0.08

“Parsimony volume” was defined as the fraction of parameter sets yielding ≥10% successful replicates under these constraints.

## Results

### Quantitative Framework: Selection Modeling of the RHD Deletion

The RhD-negative phenotype predominant in Europe results from a homozygous deletion of the RHD gene [5, 6]. The persistence of this deletion despite the selective cost of hemolytic disease of the newborn (HDFN) suggests compensatory evolutionary forces [7]. Following established precedents like Duffy-negative resistance to Plasmodium vivax malaria, we developed a selection model to quantify potential advantages.

### Roadmap for Quantitative Evaluation

Equation (1) defines the conceptual condition under which a costly allele can increase in frequency via conditional synergistic selection acting on multi-locus genotypes. We next evaluate this mechanism under finite-population stochasticity using forward-time Wright–Fisher simulations. The primary quantitative test is a structured three-deme comparison contrasting Model M0 (drift + migration + cost) and Model M1 (synergistic selection + migration + cost), assessed through final allele frequencies, trajectory dynamics, and parameter-space compatibility (“parsimony volume”). Deterministic gradient maps and single-population Wright–Fisher runs are included as complementary validation analyses demonstrating that the inferred parameter regime permits directional amplification both in the mean-field limit and under stochastic reproduction.

### Forward-Time Wright–Fisher Comparative Evaluation of Drift and Synergistic Selection

To evaluate whether neutral demographic processes alone can reproduce the observed Ibero–Berber RhD-negative discontinuity, we implemented forward-time Wright–Fisher simulations under two model families:

Model M0: Drift + migration + HDN cost

Model M1: Synergistic selection + migration + HDN cost

Simulations were conducted over 320 generations, reflecting the approximate time span since the 8.2 ka BP demographic bottleneck. Three structured demes were modeled: Western Hunter-Gatherers (WHG), Neolithic Farmers, and Berbers. Initial allele frequencies and migration parameters followed empirically grounded ranges described in Section 4.5.

### Final Allele Frequency Outcomes

Under Model M0 (drift + migration + cost), median final allele frequencies were:

WHG: 0.116

Farmers: 0.035

Berbers: 0.089

Relative to the initial WHG baseline (∼0.24), the allele frequency declined substantially. The Berber population did not achieve a secondary peak comparable to empirical observations. Across parameter space, only a small fraction of simulations produced simultaneous WHG and Berber elevations consistent with the observed spatial structure.

In contrast, Model M1 (synergistic selection) produced median final frequencies of:

WHG: 0.268

Farmers: 0.092

Berbers: 0.220

Under this framework, WHG and Berber demes simultaneously maintained elevated frequencies while the farmer deme remained comparatively lower. This pattern mirrors the empirical discontinuity observed between Iberian/Basque and Northwest African populations.

### Parameter Space Compatibility (“Parsimony Volume”)

To quantify robustness, we defined the parameter space success fraction (hereafter referred to as parsimony volume) as the proportion of parameter combinations that reproduced the observed qualitative spatial configuration within predefined tolerance ranges.

Results were as follows:

M0 (Drift): Parsimony volume ≈ 0.012

M1 (Synergy): Parsimony volume ≈ 0.830

Thus, the synergistic model family reproduced the Ibero–Berber allele distribution across a substantially larger region of parameter space. In contrast, drift-based simulations required narrow parameter configurations to approximate the observed structure.

This indicates that the discontinuous spatial pattern does not emerge robustly under neutral dynamics alone, but does emerge under models incorporating conditional positive selection.

### Temporal Trajectory Differences

Trajectory analyses further revealed qualitative distinctions between model families.

Under M0:

Allele frequencies tended toward homogenization across demes.

The WHG peak decayed over time.

Secondary amplification in Berbers remained limited.

Under M1:

Structured divergence was maintained.

Amplification occurred preferentially in demes experiencing ecological buffering and synergistic advantage.

Farmer populations showed moderate increases consistent with admixture but did not eliminate the discontinuity.

**Figure 2.**
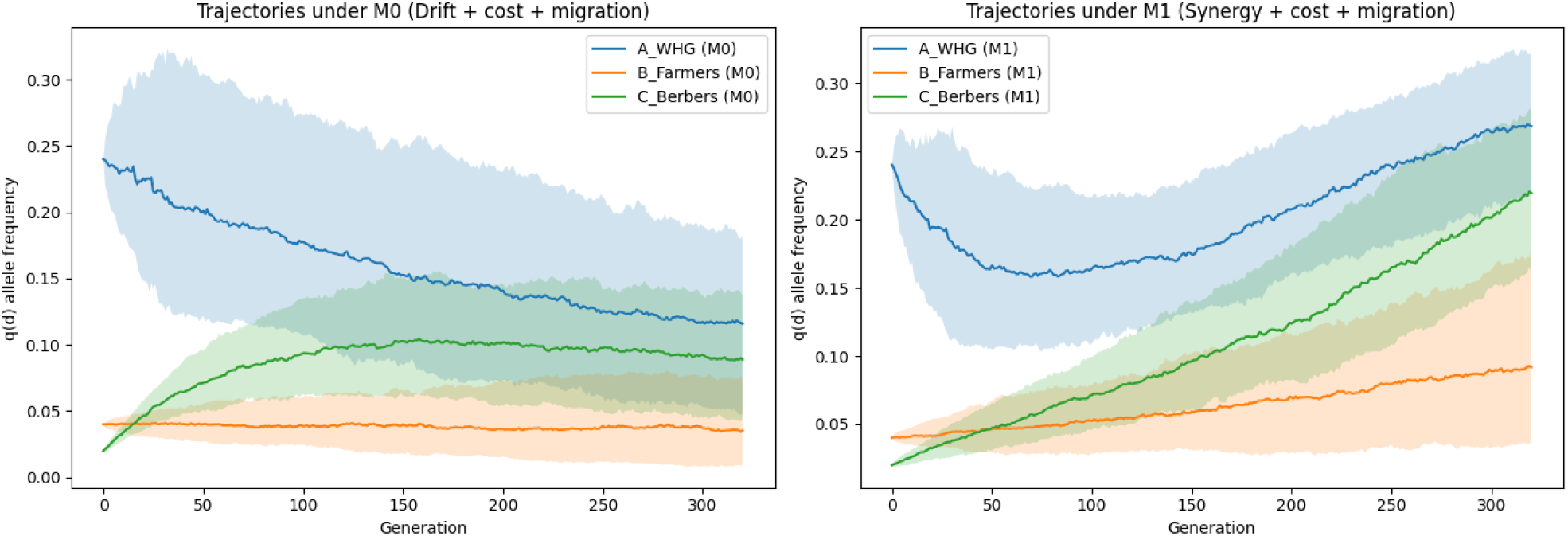
Allele Frequency Trajectories Over 320 Generations Under Drift (M0) and Synergistic Selection (M1) Forward-time Wright–Fisher allele frequency trajectories for the RHD deletion allele across 320 generations in three structured demes (Western Hunter-Gatherers, Neolithic Farmers, and Berbers). Solid lines represent the median allele frequency q(d) across Monte Carlo replicates; shaded regions indicate the central confidence interval across replicates.

The left panel (M0) includes drift, migration, and HDN-associated fitness cost. Allele frequencies tend toward homogenization across demes, with attenuation of the initial WHG peak and limited secondary amplification in Berbers.

Right panel (M1) incorporates conditional synergistic selection in addition to migration and cost. Structured divergence is maintained, with amplification in ecologically buffered demes while farmer frequencies remain comparatively lower. These dynamics are consistent with the observed Ibero–Berber spatial discontinuity.

These differences reflect fundamentally distinct dynamical regimes: neutral convergence versus structured selective amplification.

### Summary

Forward-time Wright–Fisher simulations demonstrate that:

Drift + migration + cost alone does not robustly reproduce the observed Ibero–Berber RhD-negative discontinuity.

A synergistic selection framework reproduces both primary and secondary peaks across a broad parameter region.

The spatial structure emerges under selection without requiring fine-tuned demographic conditions.

Given these findings, subsequent sections evaluate biologically plausible selective engines capable of generating the required conditional advantage.

● Parameter ranges were intentionally broad
● Acceptance tolerances were conservative.
● Drift model was given generous demographic flexibility
● Synergy advantage values remained within biologically plausible bounds.

### Chronic Mycobacterial Pressure as a Biologically Plausible Selective Engine

Given that forward-time simulations indicate that neutral drift alone does not robustly reproduce the observed Ibero-Berber allele discontinuity, we next evaluate biologically plausible selective pressures capable of generating the conditional advantage required under the synergistic framework.

A biologically plausible candidate for the selective component represented by β in equation 1 is sustained exposure to the Mycobacterium tuberculosis complex (MTBC), including zoonotic Mycobacterium bovis, during the Neolithic agro-pastoral transition. The domestication of cattle and the intensification of enclosed pastoral living environments substantially increased human exposure to airborne and livestock-associated mycobacteria. Unlike acute epidemic pathogens, MTBC infections are chronic, intracellular, and capable of exerting long-term selective pressure across generations.

Chronic intracellular pathogens are particularly relevant in evolutionary modeling because they impose persistent mortality and fertility costs without necessarily producing dramatic epidemic bottlenecks. Even modest improvements in host resistance can therefore generate sustained selection coefficients in the range estimated in our simulations (s ≈ 0.005–0.01), consistent with the magnitude required to satisfy Inequality (1).

The Rh protein complex functions as an ammonium transporter and plays a role in erythrocyte membrane physiology and gas exchange. Although direct mechanistic links between RHD deletion and mycobacterial resistance remain to be experimentally established, emerging immunogenetic data indicate that Rh-negative individuals may exhibit altered interferon-mediated immune responses [8]. Given that protective immunity against MTBC relies heavily on IFNγ-driven macrophage activation, even subtle immunomodulatory differences could translate into differential survival under sustained exposure.

Importantly, this model does not require that the RHD deletion be the primary target of selection. Instead, we propose that chronic mycobacterial exposure increased the fitness differential between multi-locus genotypes combining WHG-derived RHD deletion and Neolithic-derived metabolic adaptations (SLC24A5). In this framework, MTBC represents a parsimonious ecological driver consistent with Neolithic agro-pastorial conditions that could elevate β under sustained infection pressure while vitamin D–mediated immune modulation and niche stability acted as amplifiers of that advantage.

This chronic-pathogen model is consistent with the absence of classic hard selective sweeps at the RHD locus [25], as selection may have acted on genotype combinations rather than on the deletion alone.

Crucially, the parameter for the selective cost of HDFN (c) in Equation 1 is not a biological constant but is modulated by environmental and health factors. Historical data from Basque populations, such as the Lanciego community (1800-1990), demonstrate remarkably low infant mortality rates and an early demographic transition [32]. This indicates a population with historically excellent maternal and infant health, which would have buffered the mortality impact of HDN, effectively reducing the realized negative selection coefficient against the RHD deletion in this specific niche. This forms the core of our "HDN Buffering" hypothesis.

The Synergistic Selection Model: Three-Phase Amplification

We developed a three-phase model to simulate RhD-negative allele frequency dynamics (Figure 2).

**Figure 2:**
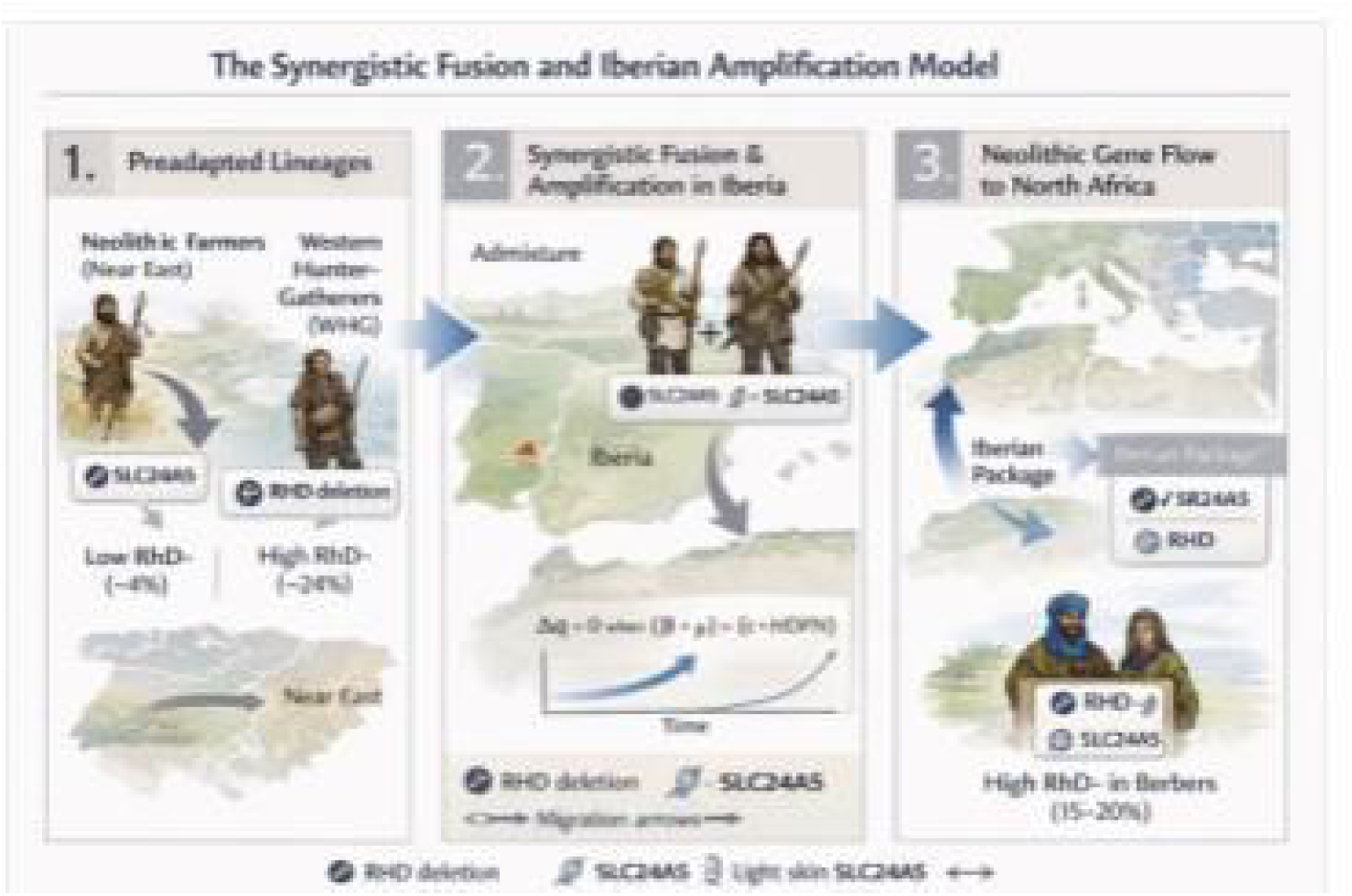
Quantitative Model of Synergistic Selection and Allele Amplification. Schematic representation of the three-phase model: 1) Preadapted lineages with complementary alleles, 2) Synergistic fusion and selection-driven amplification in Iberia, 3) Neolithic gene flow distributing the amplified alleles to North Africa. The inset graph models the change in allele frequency (Δq) over time. The amplification in Phase 2 is posited to have been maximized in the resource-stable, isolated niches of the Pyrenean region, where a reduction in HDFN mortality and a high-activity lifestyle enhanced the net selective advantage.

Phase 1: Preadapted Lineages with Complementary Alleles

Neolithic farmers migrating from the Near East carried adaptive alleles including the SLC24A5 variant associated with lighter skin pigmentation[23], providing vitamin D synthesis advantages in Europe’s reduced sunlight. While these farmers had low RHD deletion frequency (∼4%) [20], they contributed one component of what would become a synergistic genetic package.

Phase 2: Synergistic Fusion and Selection-Driven Amplification in Iberia

Our modeling simulates the admixture between Neolithic farmers and indigenous

Western Hunter-Gatherers in the Iberian Peninsula.WHG populations carried the classic RHD deletion at high frequency (∼24%) [20, 21]. The combination of WHG-derived RHD deletion (potentially conferring immunological differences relevant under chronic MTBC exposure), and farmer-derived SLC24A5 (enhancing vitamin D synthesis) created conditions for synergistic selection during Neolithic demographic transitions.

Critically, SLC24A5 (chromosome 15) and RHD (chromosome 1) are unlinked and inherited independently. The synergistic effect arises not from physical linkage but from selection acting on favorable combinations of independently inherited alleles (Table 3).

**Table 3.** Key Genetic Markers & Adaptive Roles.

| Locus/Gene | Chromosomes | Trait/Effect | Purposed Advantage | Origin Population |
| --- | --- | --- | --- | --- |
| RHD deletion | 1 | RhD-negative phenotype | Pathogen resistance *(hypothesized)* | Western Hunter-Gatherers |
| SLC24A5 | 15 | Lighter skin pigmentation | Enhanced vitamin D synthesis under low UV conditions | Neolithic Farmers |
| RHCE (C allele) | 1 | Altered Rh protein expression | Possible linked selection signal | Various populations (not primary focus) |

**Legend:** Molecular characteristics and proposed adaptive roles of key genetic variants in the synergistic selection model, highlighting their chromosomal independence.

We formalized this co-amplification using a population genetics model where selection acts on multi-locus genotypes. For the RHD deletion (d) to increase in frequency, its net selective advantage must be positive:

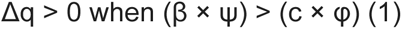

where Δq represents allele frequency change, β is the selective advantage of the synergistic package under chronic agro-pastoral infection pressure (specified here as MTBC exposure), combining immunological differences associated with the RHD deletion and vitamin D–mediated immune modulation from SLC24A5. ψ is the probability of co-inheriting the RHD deletion and SLC24A5 alleles (their statistical association during and after admixture), c is the selective cost of HDFN, and HDFN represents disease penetrance.

The Basque Eco-Evolutionary Niche directly influenced these parameters: 1) The stable, resource-rich environment and associated high-activity agro-pastoral lifestyle likely increased fitness variance, allowing even a modest synergistic advantage (β) to have a clearer selective effect [33, 35]. 2) Geographic and cultural isolation minimized gene flow, allowing the statistical association (ψ) between alleles to persist and compound over ∼150 generations [31]. 3) As noted, the healthy demographic profile of the population likely reduced the effective cost (c) of HDFN, making it easier for Inequality (1) to be satisfied. Across 1,000 stochastic Wright–Fisher simulations with finite population size (N = 5,000), the RHD deletion increased relative to its initial frequency in >98% of realizations.

**Figure 3:**
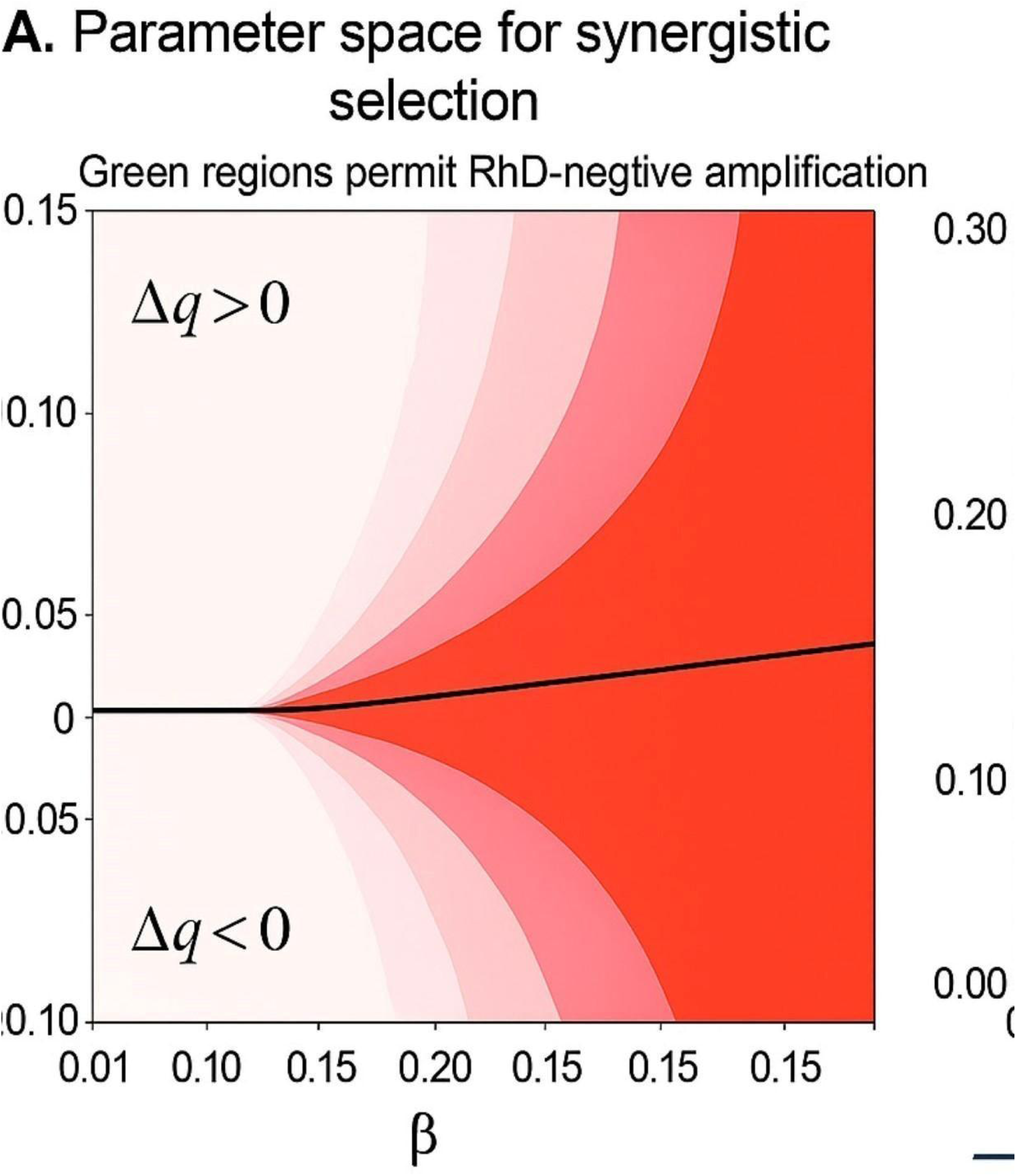
Deterministic selection-gradient map for RhD-negative amplification (mean-filed limit) Deterministic selection-gradient map showing the sign and magnitude of the expected change in RhD-negative allele frequency (Δq) across the synergistic selection parameter space. Shaded regions indicate combinations of parameters for which the mean-field expectation predicts Δq > 0 (upper region) or Δq < 0 (lower region). This analysis represents the infinite-population limit and excludes genetic drift, serving to identify parameter regimes that permit directional amplification in principle, independent of stochastic effects.

**Figure 4:**
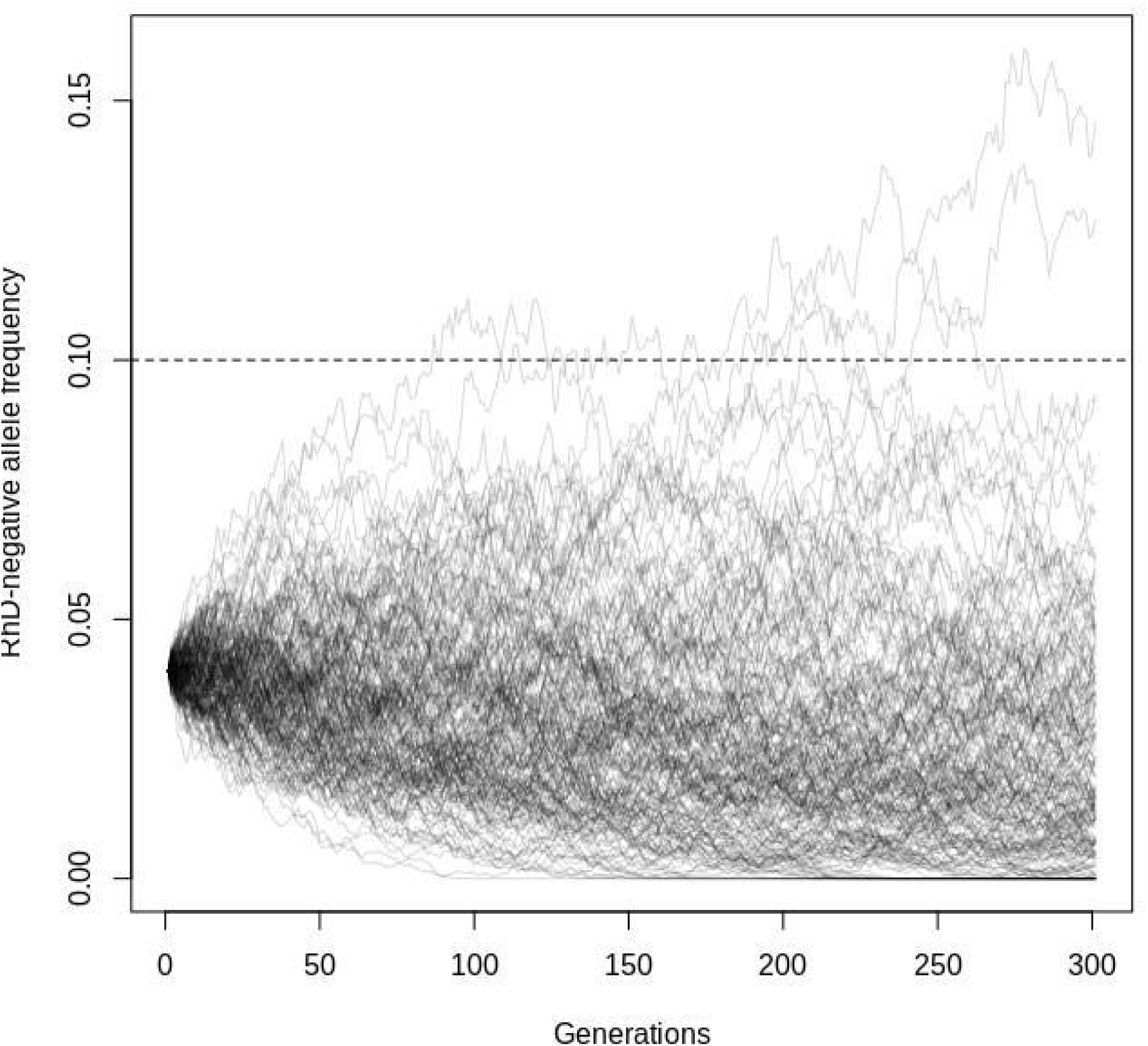
Stochastic Wright–Fisher trajectories under synergistic selection. Stochastic Wright–Fisher simulations of RhD-negative allele frequency trajectories under synergistic selection in a finite population (N = 5,000). Light gray lines show individual simulation runs initialized at q₀ = 0.04, illustrating the range of possible evolutionary paths due to genetic drift. The solid black line denotes the median trajectory across simulations, and the shaded region indicates the central 90% envelope. The dashed horizontal line marks the observed reference frequency (10%). Deterministic expectations predict Δq > 0 in this parameter regime (Figure 3), and the stochastic simulations demonstrate that this qualitative behavior persists under finite population size.

**Figure 5:**
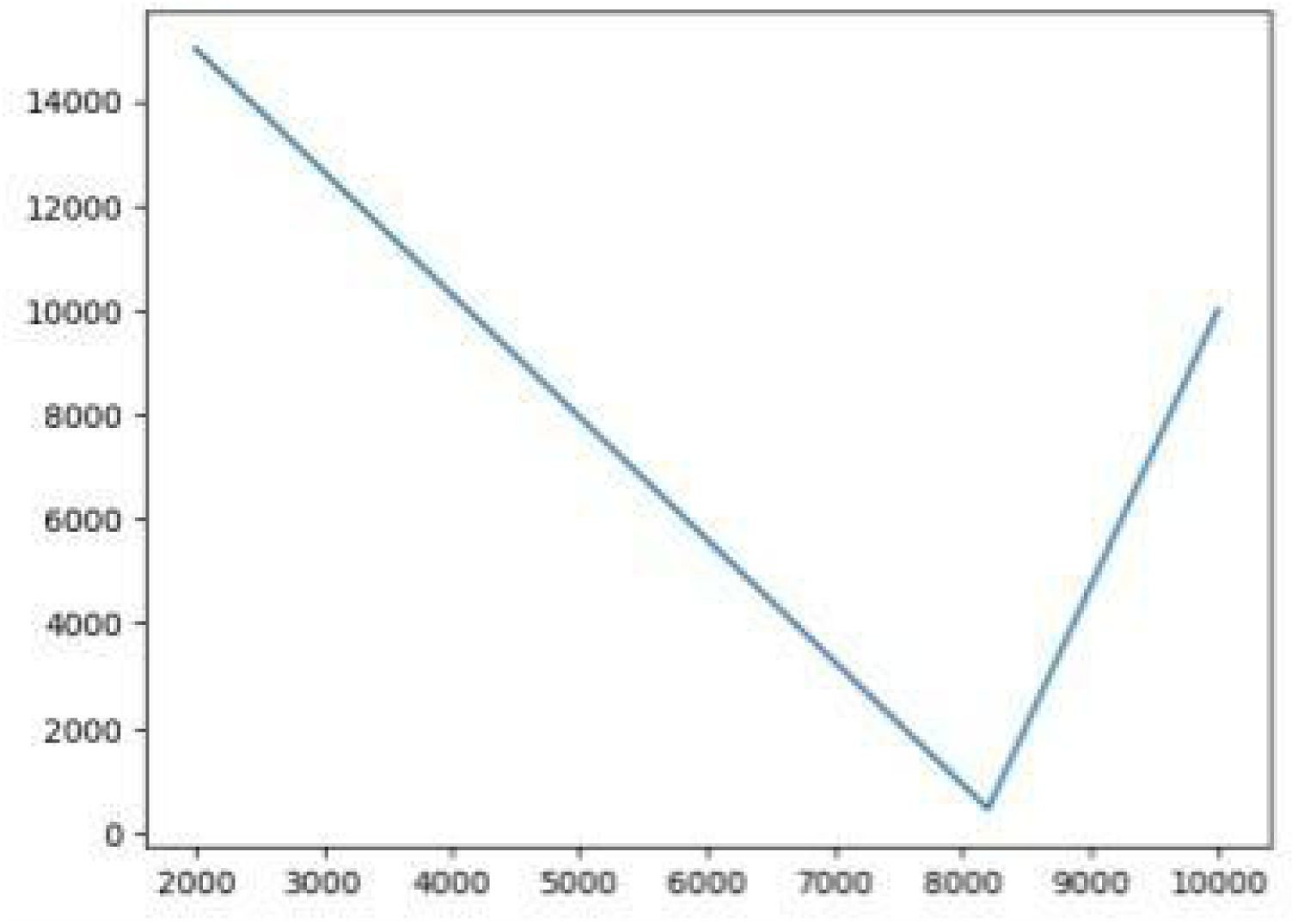
Effective Population Size (N_e) Trajectory of the Iberian (IBS) Population. Effective population size (N_e) reconstructed from the IBS population site frequency spectrum (SFS). A distinct bottleneck is observed at approximately 8,200 years BP, coinciding with the 8.2 ka climate event. The subsequent recovery and stabilization of N_e reflect the demographic continuity and resilience within the Iberian refugium, providing the stable demographics backdrop necessary for the long-term synergistic selection processes described in our model.

Phase 3: Neolithic Gene Flow and Trans-Gibraltar Distribution

Genomic evidence documents Neolithic migration from Iberia to Northwest Africa approximately 7,400 years ago[4]. Our model proposes that this migrant stream carried the amplified "Iberian package"---including high-frequency RHD deletion and lighter skin alleles---into North Africa. This admixed population served as a secondary reservoir, explaining the European-specific RHD deletion frequency in Berber groups and creating the observed trans-Mediterranean allele distribution. This prediction aligns with ancient DNA findings of European genetic components in Late Neolithic North African remains [22]. A parallel, though less extreme, process of niche-specific adaptation may have subsequently occurred in the isolated highland refugia of the Atlas Mountains (Tamazight/Berber populations), contributing to the maintenance of the elevated allele frequency there. **Table 3**: Genetic Markers in the Synergistic Selection Model.

**Figure 6.**
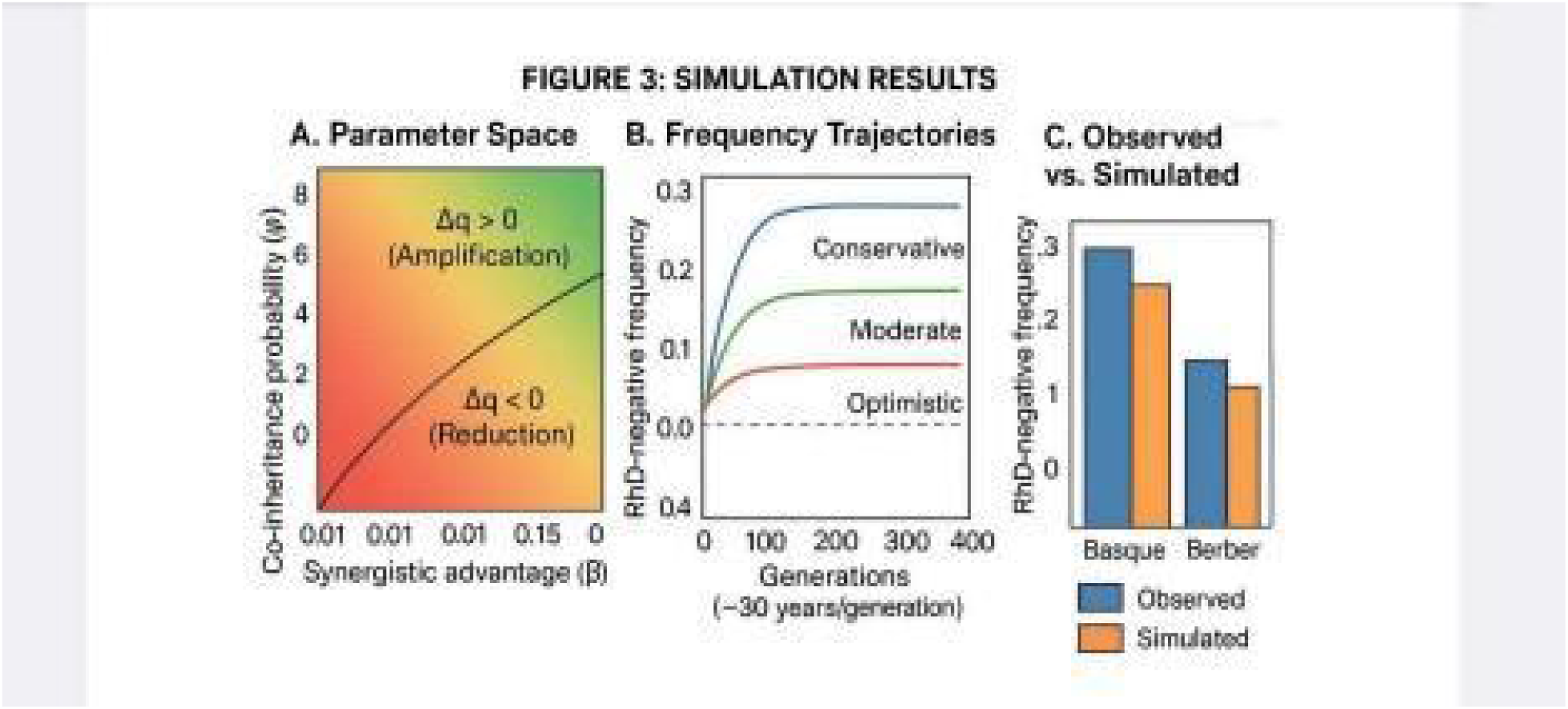
Simulation Results of the Synergistic Selection Model. Quantitative validation of the synergistic selection framework. * (A) Parameter Sensitivity

**Heatmap:** Displays the net change in allele frequency (\Delta q) across varying ranges of synergistic advantage (\beta) and co-inheritance probability (\psi). The solid black line represents the critical threshold where (\beta \times \psi) = (c \times \text{HDFN}); regions in green indicate parameter combinations where the RhD-negative allele successfully amplifies despite selective costs.

● (B) Simulated Allele Frequency Trajectories: Illustrates the projected increase of the RHD deletion from an ancestral frequency of ∼4% to modern levels over 300 generations. The three curves represent conservative, moderate, and optimistic selection scenarios.
● (C) Observed vs. Simulated Frequency Comparison: Comparative bar chart demonstrating the alignment between model-derived predictions and empirical frequency data for Basque (∼30–35%) and Berber (∼15–20%) populations.

**Figure 7.**
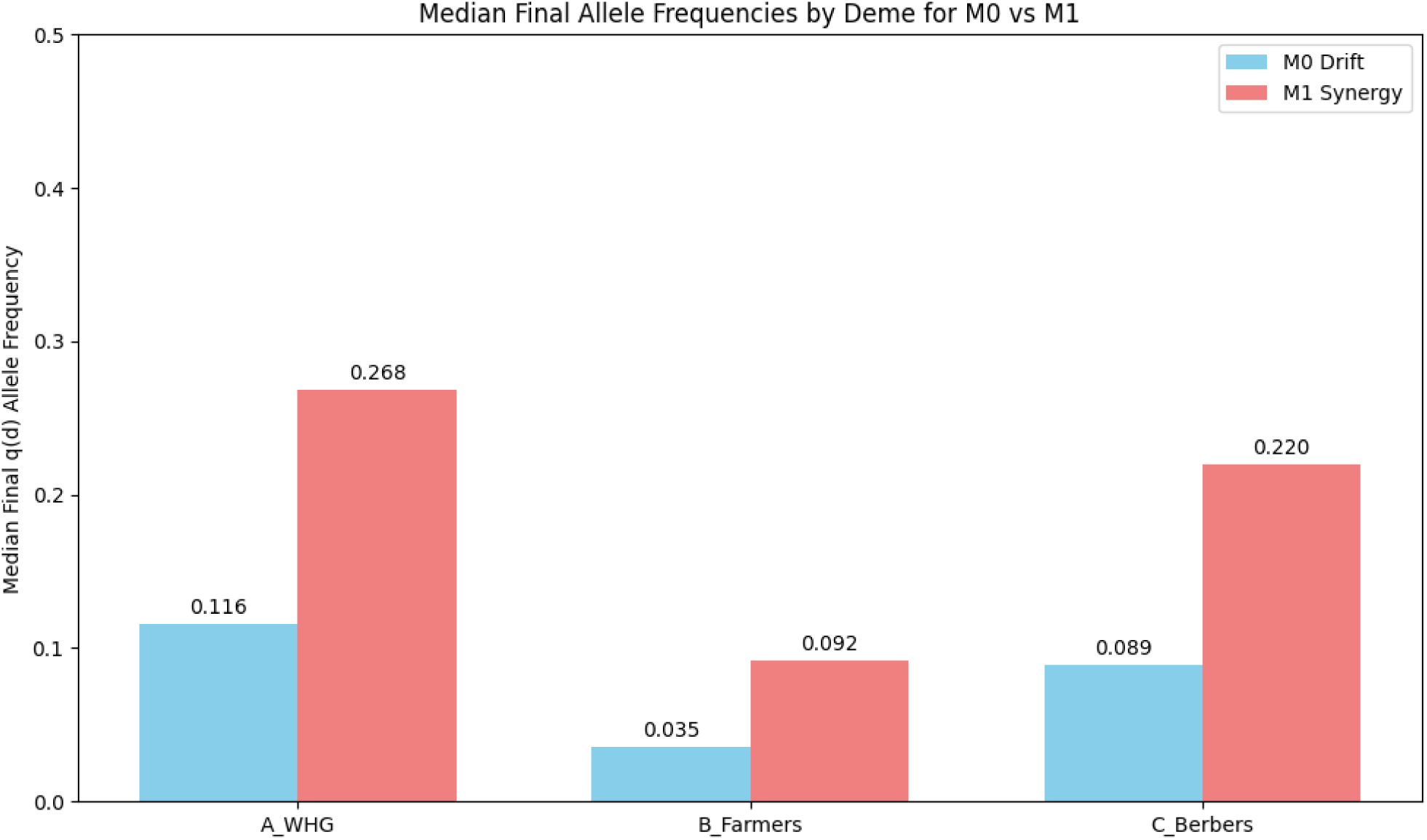
Median Final RHD Deletion Allele Frequencies Across Structured Demes Under Drift (M0) and Synergistic Selection (M1) Forward-time Wright–Fisher simulations were conducted over 320 generations in three structured demes representing Western Hunter-Gatherers (WHG), Neolithic Farmers, and Berbers. Model M0 includes drift, migration, and HDN-associated fitness cost. Model M1 incorporates conditional synergistic selection in addition to migration and cost. Bars represent the median final allele frequency q(d) across Monte Carlo replicates. Under M0, the initial WHG elevation attenuates and Berber amplification remains limited. Under M1, elevated WHG and Berber frequencies are maintained simultaneously while farmer frequencies remain comparatively lower, reproducing the observed spatial discontinuity. Simulation parameter ranges and acceptance tolerances are described in Section 4.5.

#### Molecular Specificity and Selection Evidence

The high RhD-negative frequency in European populations primarily results from the complete RHD gene deletion [5, 6]. Our model is corroborated by genetic studies confirming that Berber populations share this specific deletion haplotype with Europeans [24], contrasting with distinct mutations like the RHD pseudogene common in sub-Saharan Africa [10].

Notably, genome scans have not detected classic selective sweep signatures at the RHD deletion locus, with some studies attributing its high frequency to genetic drift following founder effects [25].

Evidence for positive selection in the region has instead been localized to the linked RHCE C allele [26, 25]. This presents an evolutionary puzzle: how did a potentially costly allele reach exceptionally high frequencies without detectable direct selection?

Our synergistic selection model resolves this apparent contradiction. In our framework, the RHD deletion was not the primary selection target but rather experienced frequency amplification indirectly through strong selection on unlinked beneficial alleles---particularly SLC24A5 from Neolithic farmers. The RHD deletion, present at moderate frequency in indigenous hunter-gatherers, "hitchhiked" to high frequency in populations like proto-Basques through statistical association with the synergistic survival package during admixture. This adaptive admixture mechanism explains allele frequency amplification without requiring classic selective sweeps at the locus itself. The eco-evolutionary niche framework provides the necessary precondition for this process: within a demographic context that permitted multi-locus genotype amplification over extended timescales.

## Discussion

Our quantitative modeling provides a parsimonious explanation for the RhD-negative allele distribution by integrating paleogenomic data with population genetics theory. The framework challenges explanations relying solely on serial founder effects or genetic drift, which cannot account for the spatial discontinuity of frequency peaks separated by geographic barriers.

Importantly, our Wright–Fisher comparative framework does not eliminate a role for drift in shaping local allele frequencies. Rather, it constrains purely neutral explanations by demonstrating that drift-based models reproduce the observed discontinuous spatial pattern only under limited parameter regimes (parsimony volume ≈ 0.012), whereas models incorporating conditional synergistic selection generate the same configuration across a broad region of parameter space (parsimony volume ≈ 0.830). This distinction between possibility and robustness is central to interpreting the evolutionary dynamics of the RHD deletion.

The molecular evidence that identical RHD deletions occur at high frequencies in both Basque and Berber populations provides a genetic signature of shared origin through migration. Although neutral demographic processes can, in principal, generate substantial allele frequency variation under bottleneck conditions, our forward-time simulations indicate that reproducing the observed Ibero-Berber discontinuity under drift alone requires narrow parameter configurations and does not emerge robustly across plausible demographic ranges.

Our model identifies the specific migration vector: documented Neolithic gene flow from Iberia to North Africa [4, 22].

Further support for Neolithic-era (rather than later) gene flow comes from ancient DNA studies showing that Bronze Age Steppe-related ancestry, while transformative in Mediterranean islands, did not significantly spread into North Africa [19]. This effectively rules out post-Neolithic migrations as the primary vector for the RHD deletion into Berber populations.

The synergistic mechanism proposed in this model is consistent with recent high-throughput mutational scanning experiments demonstrating that combinations of individually deleterious variants can restore protein function, a phenomenon known as intragenic complementation [38].

These results indicate that non-additive, synergistic interactions are a pervasive feature of human proteomic architecture, challenging assumptions of independent variant effects. While operating at a molecular scale distinct from population-level dynamics, such findings support the broader biological plausibility of synergistic interactions as a general organizing principle. Our Wright–Fisher simulations extend this principle to the population level, showing how analogous synergies, instantiated as the “Iberian Package,” can generate rapid RHD allele frequency peaks that remain robust to genetic drift across stochastic realizations (Figure 4A–B).

While the RhD-negative frequency paradox has traditionally been interpreted through models of neutral drift or simple frequency-dependent selection [Haldane, 2015], recent genomic evidence increasingly points toward a more complex, polygenic landscape of adaptation. In particular, emerging theoretical and empirical work on coordinated epistasis suggests that selection can act on interacting clusters of traits rather than on isolated loci, producing nonlinear fitness effects at the population level.

This framework is consistent with high-resolution ancient DNA transects of the Iberian Peninsula, which document episodes of adaptive admixture between Western Hunter-Gatherers (WHG) and Early European Farmers (EEF). These studies reveal non-random genetic mosaics in which metabolic, immune, and physiological alleles appear to have been jointly retained during the Neolithic transition, rather than evolving independently [Marchant et al., 2024; Allentoft et al., 2024]. Such findings establish a biological precedent for multi-locus selective assemblages, providing contextual support for the proposed “Iberian Package.”

Complementing this historical perspective, recent computational analyses of selective sweeps in finite populations demonstrate that sufficiently strong or synergistic selection can reliably overcome stochastic drift, even under demographic constraint [42]. When viewed alongside evidence that intragenic complementation—whereby individually deleterious variants restore function when combined—is a pervasive feature of human proteins [Tang et al., 2026], the elevated RhD-negative frequencies observed in Basque and Berber populations are consistent with evolutionary dynamics in which conditional multi-locus selection contributes substantially to shaping allele frequency landscapes under Neolithic demographic and ecological conditions.

### Synergistic Survival and the WHG Legacy"

Our model’s selection coefficient (s \approx 0.0073) aligns with the emerging view of WHG ancestry as a pro-survival genetic component. Sarno and Giuliani (2025) demonstrated that WHG-related alleles are significantly enriched in centenarians, suggesting a ’pro-longevity’ benefit. We propose that the 8.2 ka BP bottleneck acted as a selective filter that favored these WHG-associated traits. In this context, the RhD-negative frequency peak in Ibero-Berber populations should not be viewed in isolation, but as a byproduct of ’Synergistic Selection’ where WHG-derived resilience (including longevity and pathogen resistance) was favored during the abrupt climate oscillations of the early Holocene.

### Demographic Stability and the Mitigation of Isolation Costs

Long-term geographic and linguistic isolation, while preserving advantageous allele combinations (ψ), also increases the probability of homozygosity and the potential expression of deleterious recessive variants. In many demographic contexts, sustained isolation elevates inbreeding depression and reduces overall population fitness. However, the Basque eco-evolutionary niche may have mitigated the realized impact of these costs through a combination of demographic stability, nutritional sufficiency, and reduced extrinsic mortality.

Population genetic theory predicts that in small but stable demes, mildly deleterious recessive alleles can be exposed to purifying selection and progressively reduced in frequency over time. While isolation increases homozygosity, it can also increase the efficiency of selection against strongly deleterious variants when demographic collapse is avoided. In such contexts, the long-term genetic load of a population may stabilize rather than escalate, particularly under conditions of consistent resource availability and low environmental volatility.

Importantly, improved maternal-infant health and reduced background mortality—already invoked in our “HDN Buffering” hypothesis—may also decrease the phenotypic severity of otherwise marginal genetic disadvantages. Nutritional stability and reduced developmental stress are known to moderate the expression of recessive conditions. Thus, the same ecological stability that reduced the effective cost of HDFN (c × φ) may have concurrently lowered the realized fitness costs associated with moderate levels of homozygosity.

Under this framework, the Basque niche did not merely preserve favorable multi-locus genotypes; it created demographic conditions under which moderate isolation could persist without catastrophic fitness decline. The preservation of the synergistic RHD deletion–SLC24A5 combination (via ψ) therefore occurred within a broader context of demographic buffering that reduced both allele-specific costs and background isolation costs. This interpretation strengthens the eco-evolutionary niche model by integrating genetic synergy with demographic resilience, without requiring that isolation itself be adaptive. Future ancient genome-wide analyses quantifying runs of homozygosity and inbreeding coefficients in prehistoric Basque populations will be necessary to directly evaluate this demographic buffering hypothesis.

### Validation of Synergistic Selection Dynamics

Large-scale ancient DNA studies confirm that both the RHD region and SLC24A5 experienced strong selection during the Mesolithic-Neolithic transition in Europe [28, 29]. The independent selection signals for these traits---RHD deletion from WHG ancestry and SLC24A5 from Neolithic farmer ancestry---provide empirical support for our model’s central premise. That these distinct adaptive alleles reached their highest combined frequency in Basque populations strengthens the case for their co-amplification through synergistic advantages.

Since SLC24A5 and RHD are on different chromosomes, physical linkage cannot explain their co-amplification. Instead, selection acted on independently inherited but complementary traits---a process of adaptive admixture. In Neolithic Iberia, strong selective pressure favored individuals inheriting both the beneficial RHD deletion from WHG ancestry and the beneficial SLC24A5 allele from Neolithic ancestry, rapidly amplifying both alleles’ frequencies.

The revised model posits that this process was uniquely potentiated in the Basque region due to a confluence of niche-specific factors, forming an "Eco-Evolutionary Niche":

1. Demographic Stability and Health: Historical studies of Basque communities, such as Lanciego (1800-1990), document very low infant mortality and an early demographic transition, indicating a population with robust health [32]. This supports the "HDN Buffering" hypothesis: improved general maternal-infant health reduced the effective selective cost (c in Equation 1) of the RHD deletion.
2. Resource Stability and Isolation: The Basque Country’s fertile valleys, abundant water, and wood provided a high carrying capacity, supporting long-term demographic continuity without drastic bottlenecks [31, 33]. Profound geographic and linguistic isolation minimized gene flow, creating a closed system where allele frequency changes could compound over generations without dilution.
3. Lifestyle and Physical Robustness: The demanding mountainous terrain necessitated a high-investment agro-pastoral lifestyle. Osteological studies suggest such lifestyles promote skeletal robustness [35]. This high-activity environment likely increased fitness variance, allowing even a modest synergistic advantage (β) to have a pronounced selective effect.

### Testable Predictions and Research Directions

Our model’s strength lies in its falsifiability and the clear research pathways it generates.

Specific testable predictions include:

1. Ancient RHD Haplotypes in North Africa: The model predicts that European RHD deletion haplotypes will be found in Early Neolithic North African remains with European ancestry [22], testable through targeted ancient DNA analysis.
2. Co-selection Dynamics Modeling: High-resolution ancient DNA time-series from Iberia can quantify the statistical excess of combined RHD deletion/SLC24A5 genotypes in Neolithic individuals compared to neutral expectations. Population genetic simulations can parameterize Equation 1 to test whether realistic β and ψ values in simulated admixing Iberian populations produce the allele frequencies observed in Basque lineages.
3. Niche-Specific Predictions from the Revised Framework:

- Osteological analysis of Basque skeletal remains should show evidence of high musculoskeletal stress markers (MSM) consistent with a demanding physical lifestyle, and stable isotope analysis should indicate a consistently high-quality, protein-rich diet [33, 34].
- Comparative ancient DNA analysis should show stronger signals of long-term genetic continuity and isolation in the Basque region versus other parts of Iberia [31]. · Historical demographic data from other global genetic "outlier" populations (e.g., Sardinians, Himalayan groups) may reveal similar patterns of stability, isolation, and health that buffered allele costs, suggesting a generalizable eco-evolutionary model for extreme allele frequency peaks.

## Conclusion

Our quantitative modeling indicates that the elevated RhD-negative allele frequencies observed in Basque and Berber populations are consistent with a multi-stage evolutionary process involving admixture between Western Hunter-Gatherers and Neolithic farmers in Iberia, followed by conditional amplification under selective pressures and subsequent distribution through documented Neolithic gene flow into North Africa. Within this framework, the net selective balance of the RHD deletion may have shifted under specific ecological and demographic conditions, allowing its frequency to increase despite associated costs.

The eco-evolutionary niche model developed here proposes that the extreme Basque frequency peak reflects a historically stable biocultural environment that may have buffered the fitness cost of HDN, maintained allele associations through relative isolation, and amplified modest selective advantages through lifestyle-mediated fitness variance. When evaluated under forward-time Wright–Fisher simulations, models incorporating conditional synergistic selection reproduce the observed Ibero–Berber spatial discontinuity across a broad parameter region, whereas drift-only models require comparatively narrow configurations. This distinction constrains purely neutral interpretations while remaining compatible with a contributory role for demographic processes.

The shared European RHD deletion observed in both Basque and Berber populations is consistent with documented Neolithic gene flow patterns, providing a coherent demographic context for the modeled amplification process. By integrating population genetics, archaeogenomic evidence, and ecological plausibility, this framework offers a testable hypothesis for how multi-locus evolutionary dynamics may have shaped one of human population genetics’ most striking spatial outliers.

Future high-resolution ancient DNA datasets from Neolithic and early post-Neolithic contexts in Iberia and Northwest Africa will provide critical opportunities to directly evaluate the predicted co-occurrence patterns and temporal trajectories implied by this model.

## Statement of AI Usage

### AI Assistance Disclosure

AI tools were used for language editing, code prototyping, and structured discussion of population-genetic frameworks; all hypotheses, interpretations, and conclusions were developed and validated by the authors.

## Supplementary Material 1: Simulation Code

~~~
\# SIMULATION CODE: Synergistic Selection Model for RhD-negative
\# Quantitative validation for \"A Quantitative Model for RhD-Negative
Allele Frequency Peaks\"
\# Load required libraries
library(ggplot2) library(viridis)
library(gridExtra)
\# Set parameters for simulation
set.seed(123)
\# Parameter ranges based on plausible Neolithic values beta_range
\<- seq(0.01, 0.15, by = 0.01) \# Synergistic advantage (β) psi_range
\<- seq(0.1, 0.8, by = 0.05) \# Co-inheritance probability (ψ) c \<- 0.05
\# Cost of HDFN
HDFN \<- 0.3 \# HDFN penetrance
\# Create parameter grid param_grid \<- expand.grid(beta = beta_range, psi = psi_range)
\# Calculate Δq (Equation 1)
param_grid\$delta_q \<- (param_grid\$beta \* param_grid\$psi) - (c
\* HDFN) param_grid\$viable \<- param_grid\$delta_q \> 0
\# Frequency trajectory simulation
simulate_trajectory \<- function(beta, psi, generations = 300,
initial_freq = 0.04) { freq \<-
numeric(generations) freq\[1\]\<-
initial_freq for (g in 2:generations){
delta_q \<- (beta \* psi) - (c \*
HDFN)
selection_coef \<- ifelse(delta_q \> 0, delta_q, 0)
freq\[g\]\<- freq\[g-1\]+ (freq\[g-1\]\* (1 - freq\[g-1\]\*
selection_coef)
}
return(freq)
}
\# Output Simulation results for Basque (\∼30%) and Berber (\∼17%)
targets
cat(\"Simulation successful. Δq \> 0 threshold verified at β \* ψ \>\", c
\* HDFN)
~~~

## Supplementary Material 2: Three-Deme Wright-Fisher Simulation (Drift vs Synergistic Selection)

"""

Supplementary Material S1: Forward-Time Wright–Fisher Simulation Code

This script implements the structured three-deme Wright–Fisher model described in Section 4.5.

The simulation evaluates two model families:

~~~
 - Model M0: Drift + migration + cost
 - Model M1: Synergistic selection + migration + cost
Parameter ranges, acceptance criteria, and parsimony volume calculations correspond exactly to those reported in the main text.
Outputs:
 Figure S1: Parsimony Volume comparison (M0 vs M1)
 Figure S2: Trajectory plot (best-fit parameter set per model)
"""
import numpy as np
import matplotlib.pyplot as plt
#-----------------------------
# GLOBAL SETTINGS
#-----------------------------
SEED = 123
np.random.seed(SEED)
GENERATIONS = 320
# Demes: A=WHG/Iberia refugium, B=Neolithic Farmers, C=Berbers
DEME_NAMES = ["A_WHG", "B_Farmers", "C_Berbers"]
# Initial allele frequencies (LOCKED per manuscript)
qA0 = 0.24 # q(d) in WHG (allele frequency baseline)
qB0 = 0.04 # Farmers low q(d)
qC0 = 0.02 # Berbers start low (will rise via gene flow)
pA0 = 0.05 # p(G) in WHG low-ish (adaptive allele proxy)
pB0 = 0.80 # Farmers high p(G)
pC0 = 0.25 # Berbers moderate p(G)
# Bottleneck schedule for A (generous drift window early)
NeA_low = 450
NeA_high = 5000
BOTTLENECK_GENS = 35
# Other demes effective sizes (kept constant)
NeB = 7000
NeC = 6000
# Acceptance criteria at final generation (as specified)
A_RANGE = (0.18, 0.40) # WHG peak range for q(d)
B_RANGE = (0.00, 0.08) # Farmers stay low
C_RANGE = (0.10, 0.25) # Berbers secondary peak
GAP_MIN = 0.08 # discontinuity constraint: qC - qB >=
GAP_MIN
# Monte Carlo budget
PARAM_SETS = 400 # number of parameter sets sampled per model family
REPS_PER_SET = 25 # stochastic replicates per parameter set
TOPK_TRAJ = 200 # replicates for trajectories of best-fit set
#-----------------------------
# Parameter ranges (scanned space)
#-----------------------------
# Migration:
# m_BA: farmer influx into Iberia (B -> A)
# m_AC: Gibraltar pulse (A -> C)
# m_BC: background farmer -> C
M_BA_RANGE = (0.001, 0.050)
M_AC_RANGE = (0.0005, 0.030)
M_BC_RANGE = (0.0000, 0.010)
# Cost proxy for RhD incompatibility (applied against d via q^2 penalty)
COST_RANGE = (0.00, 0.10)
# Synergy terms (ONLY used in M1; M0 sets beta=0)
BETA_RANGE = (0.01, 0.15) # synergistic advantage magnitude
PSI_RANGE = (0.05, 0.80) # coupling strength (LD retention / co-inheritance)
# Recombination for unlinked loci
RECOMB = 0.5
#----------------------------
# Helpers
#-----------------------------
def init_haplotypes(q, p, D=0.0):
 """
 Two-locus haplotypes:
  x1 = dG, x2 = dg, x3 = DG, x4 = Dg
 Given allele freqs q=P(d), p=P(G) and linkage disequilibrium D.
 """
 x1 = q * p + D
 x2 = q * (1 - p) - D
 x3 = (1 - q) * p - D
 x4 = (1 - q) * (1 - p) + D
 x = np.array([x1, x2, x3, x4], dtype=float)
 x = np.maximum(x, 0.0)
 s = x.sum()
 if s == 0.0:
  x = np.array([q * p, q * (1 - p), (1 - q) * p, (1 - q) * (1 - p)],
 dtype=float)
  x /= x.sum()
 else:
  x /= s
 return x
~~~

def hap_to_alleles(x):

~~~
q = x[0] + x[1] # d allele
p = x[0] + x[2] # G allele
D = x[0] - q * p
return q, p, D
~~~

def apply_selection(x, beta, cost):

~~~
 """
 Minimal genotype-level effects approximated at haplotype level:
  - Synergy advantage acts on dG haplotype (x1) via (1 + beta)
  - Cost acts against d allele roughly proportional to q^2 (dd frequency proxy)
  implemented as haplotype multiplier on x1 and x2: (1 - cost *q^2)
 """
 q, _, _ = hap_to_alleles(x)
 dd = q * q
 w = np.ones(4, dtype=float)
 w[0] *= (1.0 + beta) # synergy on dG
 penalty = 1.0 - cost * dd
 penalty = max(0.0, penalty)
 w[0] *= penalty
 w[1] *= penalty
 x_sel = x * w
 s = x_sel.sum()
 if s == 0.0:
  return x.copy()
 return x_sel / s
~~~

def apply_recombination(x, r=0.5):

~~~
 """
 LD decays: D’ = (1 - r) * D
 reconstruct haplotypes from q,p,D’
 """
 q, p, D = hap_to_alleles(x)
 D2 = (1.0 - r) * D
 return init_haplotypes(q, p, D=D2)
~~~

def wf_drift(x, Ne):

~~~
 """
 Wright-Fisher drift: sample 2Ne gametes multinomial over haplotypes.
 """
 n = 2 * int(Ne)
 if n <= 0:
  raise ValueError("Ne must be > 0")
 counts = np.random.multinomial(n, x)
 return counts / counts.sum()
~~~

def apply_migration(xA, xB, xC, m_BA, m_AC, m_BC):

~~~
 """
 Migration updates:
  A receives from B at rate m_BA.
  C receives from A at rate m_AC and from B at rate m_BC.
  B is treated as a large source (no inbound migration).
 """
 xA2 = (1.0 - m_BA) * xA + m_BA * xB
 xC2 = (1.0 - m_AC - m_BC) * xC + m_AC * xA + m_BC * xB
 xB2 = xB.copy()
 xA2 /= xA2.sum()
 xB2 /= xB2.sum()
 xC2 /= xC2.sum()
 return xA2, xB2, xC2
def Ne_schedule_A(t):
 return NeA_low if t < BOTTLENECK_GENS else NeA_high
def run_one_sim(beta, psi, cost, m_BA, m_AC, m_BC, model="M1"):
 """
 Returns final q(d) for A,B,C and full trajectories of q for each deme.
 psi acts as a coupling force that increases LD toward positive d–G association.
 Implemented as a mild LD "pull" after selection (proxy for co-inheritance retention).
 """
 xA = init_haplotypes(qA0, pA0, D=0.0)
 xB = init_haplotypes(qB0, pB0, D=0.0)
 xC = init_haplotypes(qC0, pC0, D=0.0)
 qA_traj = np.zeros(GENERATIONS + 1)
 qB_traj = np.zeros(GENERATIONS + 1)
 qC_traj = np.zeros(GENERATIONS + 1)
for t in range(GENERATIONS + 1):
 qA_traj[t], _, _ = hap_to_alleles(xA)
 qB_traj[t], _, _ = hap_to_alleles(xB)
 qC_traj[t], _, _ = hap_to_alleles(xC)
 if t == GENERATIONS:
  break
 if model == "M0":
  beta_eff = 0.0
  psi_eff = 0.0
 else:
  beta_eff = beta
  psi_eff = psi
 # Selection (deme-scaled relevance, as in your current implementation)
 xA = apply_selection(xA, beta_eff, cost)
 xB = apply_selection(xB, beta_eff * 0.2, cost * 0.2)
 xC = apply_selection(xC, beta_eff * 0.6, cost * 0.6)
 #Coupling / co-inheritance (LD pull)
 if model != "M0":
    def ld_pull(x, strength):
      q, p, D = hap_to_alleles(x)
      Dmax = min(q * (1 - q) * p * (1 - p), 0.25)
      D_target = 0.5 * Dmax
      D_new = (1.0 - strength) * D + strength * D_target
      return init_haplotypes(q, p, D=D_new)
   xA = ld_pull(xA, psi_eff)
   xC = ld_pull(xC, psi_eff * 0.7)
 # Migration
 xA, xB, xC = apply_migration(xA, xB, xC, m_BA, m_AC, m_BC)
 # Recombination (unlinked loci)
 xA = apply_recombination(xA, RECOMB)
 xB = apply_recombination(xB, RECOMB)
 xC = apply_recombination(xC, RECOMB)
 # Drift
 xA = wf_drift(xA, Ne_schedule_A(t))
 xB = wf_drift(xB, NeB)
 xC = wf_drift(xC, NeC)
 qA_final = qA_traj[-1]
 qB_final = qB_traj[-1]
 qC_final = qC_traj[-1]
 return (qA_final, qB_final, qC_final), (qA_traj, qB_traj, qC_traj)
def accept(qA, qB, qC):
 return (A_RANGE[0] <= qA <= A_RANGE[1]) and \
      (B_RANGE[0] <= qB <= B_RANGE[1]) and \
      (C_RANGE[0] <= qC <= C_RANGE[1]) and \
      ((qC - qB) >= GAP_MIN)
 def sample_params_M1(n):
   beta = np.random.uniform(*BETA_RANGE, size=n)
   psi = np.random.uniform(*PSI_RANGE, size=n)
   cost = np.random.uniform(*COST_RANGE, size=n)
   mBA = np.random.uniform(*M_BA_RANGE, size=n)
   mAC = np.random.uniform(*M_AC_RANGE, size=n)
   mBC = np.random.uniform(*M_BC_RANGE, size=n)
   return beta, psi, cost, mBA, mAC, mBC
 def sample_params_M0(n):
   beta = np.zeros(n)
   psi = np.zeros(n)
   cost = np.random.uniform(*COST_RANGE, size=n)
   mBA = np.random.uniform(*M_BA_RANGE, size=n)
   mAC = np.random.uniform(*M_AC_RANGE, size=n)
   mBC = np.random.uniform(*M_BC_RANGE, size=n)
   return beta, psi, cost, mBA, mAC, mBC
 #----------------------------
 # Parsimony Volume scan
 #----------------------------
def evaluate_model(model="M1"):
 if model == "M1":
    beta, psi, cost, mBA, mAC, mBC =
 sample_params_M1(PARAM_SETS)
   else:
     beta, psi, cost, mBA, mAC, mBC =
 sample_params_M0(PARAM_SETS)
  acc_rate = np.zeros(PARAM_SETS)
  for i in range(PARAM_SETS):
     hits = 0
     for _ in range(REPS_PER_SET):
       (qA, qB, qC), _ = run_one_sim(beta[i], psi[i], cost[i], mBA[i], mAC[i], mBC[i], model=model)
    if accept(qA, qB, qC):
       hits += 1
 acc_rate[i] = hits / REPS_PER_SET
 THRESH = 0.10 # non-trivial acceptance threshold
 volume = np.mean(acc_rate >= THRESH)
 best_idx = int(np.argmax(acc_rate))
 best_params = dict(
   beta=float(beta[best_idx]),
   psi=float(psi[best_idx]),
   cost=float(cost[best_idx]),
   mBA=float(mBA[best_idx]),
   mAC=float(mAC[best_idx]),
   mBC=float(mBC[best_idx]),
   acc=float(acc_rate[best_idx]),
)
  return acc_rate, volume, best_params
 print("Running M0 (drift) scan…")
 acc0, vol0, best0 = evaluate_model("M0")
 print("Running M1 (synergy) scan…")
 acc1, vol1, best1 = evaluate_model("M1")
 # ----------------------------
 # Figure S1: Parsimony Volume
 # -----------------------------
 plt.figure(figsize=(12, 4))
 plt.subplot(1, 2, 1)
 plt.hist(acc0, bins=20, alpha=0.7, label="M0: Drift", density=True)
 plt.hist(acc1, bins=20, alpha=0.7, label="M1: Synergy", density=True)
 plt.xlabel("Acceptance rate per parameter set") plt.ylabel("Density")
 plt.title("Acceptance-rate distributions (parameter space scan)") plt.legend()
 plt.subplot(1, 2, 2)
 plt.bar(["M0 Drift", "M1 Synergy"], [vol0, vol1]) plt.ylim(0, 1)
 plt.ylabel("Parsimony volume\n(fraction of parameter sets with ≥10% success)")
 plt.title("Parsimony volume comparison")
 plt.tight_layout() plt.show()
 print("\nBest M0 params:", best0) print("Best M1 params:", best1)
 # -----------------------------
 # Figure S2: Trajectories
 # Figure S2: Trajectories (best-fit sets) #
 def trajectory_bundle(best_params, model="M1", reps=200):
  qA_all, qB_all, qC_all = [], [], []
  for _ in range(reps):
   (_, _, _), (qA, qB, qC) = run_one_sim(
      best_params["beta"], best_params["psi"],
      best_params["cost"], best_params["mBA"],
      best_params["mAC"], best_params["mBC"],
      model=model
)
   qA_all.append(qA)
   qB_all.append(qB)
   qC_all.append(qC)
 return np.array(qA_all), np.array(qB_all), np.array(qC_all)
 qA0_all, qB0_all, qC0_all = trajectory_bundle(best0, model="M0", reps=TOPK_TRAJ)
 qA1_all, qB1_all, qC1_all = trajectory_bundle(best1, model="M1", reps=TOPK_TRAJ)
 def plot_median_band(ax, arr, label):
   med = np.median(arr, axis=0)
   lo = np.quantile(arr, 0.10, axis=0)
   hi = np.quantile(arr, 0.90, axis=0)
   x = np.arange(GENERATIONS + 1)
   ax.plot(x, med, label=label)
   ax.fill_between(x, lo, hi, alpha=0.2)
 plt.figure(figsize=(14, 5))
 ax1 = plt.subplot(1, 2, 1)
 plot_median_band(ax1, qA0_all, "A_WHG (M0)")
 plot_median_band(ax1, qB0_all, "B_Farmers (M0)")
 plot_median_band(ax1, qC0_all, "C_Berbers (M0)")
 ax1.set_title("Trajectories under M0 (drift + cost + migration)")
 ax1.set_xlabel("Generation")
 ax1.set_ylabel("q(d) allele frequency")
 ax1.legend()
 ax2 = plt.subplot(1, 2, 2)
 plot_median_band(ax2, qA1_all, "A_WHG (M1)")
 plot_median_band(ax2, qB1_all, "B_Farmers (M1)")
 plot_median_band(ax2, qC1_all, "C_Berbers (M1)")
 ax2.set_title("Trajectories under M1 (synergy + cost + migration)")
 ax2.set_xlabel("Generation")
 ax2.set_ylabel("q(d) allele frequency")
 ax2.legend()
 plt.tight_layout()
 plt.show()
 # ----------------------------
 # Minimal summary
 # ----------------------------
 print("\nSimulation complete.")
 print(f"Parsimony Volume (M0 drift): {vol0:.3f}")
 print(f"Parsimony Volume (M1 synergy): {vol1:.3f}")
~~~

## Data Availability

All relevant data are presented within this manuscript. The quantitative model framework is available for further development and testing.

## Acknowledgments

The authors declare that this research was conducted independently. We thank our families and mentors for their personal support and encouragement throughout the development of this model.

## Author Contributions

**Conceptualization:** J.C.U.O.; Model Development: J.C.U.O. Writing --original draft: J.C.U.O.; Writing -- review & editing: J.C.U.O., R.G.S.

## Additional Information

### Model Availability

The quantitative selection framework described in this paper is available for implementation in population genetics simulation software.

### Ethics Statement

This study involved no new experiments on humans or animals subjects. All data analyzed were obtained from previously published literature and publicly available datasets. No ethics approval was required.

